# Basal ganglia high-frequency activity is co-modulated with the phase of motor cortical beta and shifted between the subthalamic nucleus and the internal pallidum during sustained motor control

**DOI:** 10.1101/2020.08.30.273888

**Authors:** Petra Fischer, Alek Pogosyan, Alexander L. Green, Tipu Z. Aziz, Jonathan Hyam, Thomas Foltynie, Patricia Limousin, Ludvic Zrinzo, Michael Samuel, Keyoumars Ashkan, Mauro Da Lio, Mariolino De Cecco, Alessandro Luchetti, Peter Brown, Huiling Tan

## Abstract

Beta oscillations are readily observed in motor cortex and the basal ganglia, but to which extent they are functionally relevant is unclear. To understand how activity transfer between different nodes of the cortico-basal ganglia network is affected by cortical beta oscillations in different behavioural conditions, we recorded local field potentials and electroencephalography (EEG) activity in a low-force motor control task and during rest in Parkinson’s patients undergoing deep brain stimulation (DBS) surgery. The patients received DBS of either the subthalamic nucleus (STN) or the internal globus pallidus (GPi), which allowed us to investigate if STN and GPi broad-band high-frequency activity (HFA; >150 Hz) is co-modulated with the phase of motor cortical beta activity. We found significant modulation patterns in the STN and the GPi, which were inverted while patients performed the task, showing that GPi activity fluctuations likely are crafted by other inputs than the direct excitatory STN afferents. We also found that consistent STN modulation disappeared during rest, showing disengagement in this condition, while GPi modulation was maintained, again evidencing that beta-band activity fluctuations in the GPi can be relatively independent of those in the STN. The difference between HFA modulation patterns in the task and rest recordings suggests a potential functional role of beta phase-locked HFA modulation in controlling sustained contractions. Examination of HFA co-modulation patterns at different sites of the cortico-basal ganglia-thalamo-cortical network under different behavioural conditions may provide a tool with which to define the impact of beta synchronization on network communication.

## Introduction

Beta oscillations (13-30 Hz) are an abundant phenomenon in the cortico-basal ganglia-thalamo-cortical network and have been associated with a wide range of potential functions, including feedback processing (Cao and Hu, 2016; Tan et al., 2016; Torrecillos et al., 2015), communicating sensorimotor information across widespread areas (Classen et al., 1998; Kilavik et al., 2013; Rubino et al., 2006), maintaining muscle synergies (Aumann and Prut, 2015), movement inhibition (Aron et al., 2016), timing (Kononowicz et al., 2019) as well as clearing out previously held information (Schmidt et al., 2019). However, if instead of assigning broad functions, we aim to link the phenomenon of these oscillations to task-specific neural computations, such as coincidence detection or recurrent amplification (Carandini, 2012; Douglas et al., 1995), we need to understand in more detail how basal ganglia and cortical activity patterns are co-modulated.

We have recently found that when Parkinson’s patients performed a low-level force adjustment task, bursts of beta oscillations in the STN and the GPi appeared when the force had to be stabilized (Fischer et al., 2019). Motor cortical and basal ganglia beta oscillations also were phase-coupled, raising the possibility that task-related communication between these areas may be structured within cycles of beta activity. Activity in each basal ganglia nucleus is a mixture of multiple inputs but could be dominated by routing along different paths. The STN projects to the GPi via a direct excitatory connection (Nambu et al., 2000), but the GPi also receives inhibitory inputs from the GPe and from striatal medium-spiny neurons (Shink and Smith, 1995; Smith et al., 1994).

One major open question is, how does basal ganglia firing in different nuclei co-fluctuate with the phase of task-related motor cortical beta activity? In this study, we set out to answer if GPi activity, a major basal ganglia output, mirrors the fluctuations in the STN in two conditions – during motor engagement and at rest. If activity is not simply mirrored, we can assume that there is a shift in the balance between different inputs to the GPi or less direct routing of STN activity via STN➔GPe➔GPi projections. Additionally, if modulation patterns differ between conditions, we will gain insights into alterations of network dynamics that may enable different task-dependent functional roles of beta oscillations.

Because simultaneous invasive recordings of STN and GPi firing activity cannot usually be performed in humans, we have employed the following workarounds: First, we compare STN and GPi activity recorded from two separate groups but both having the same motor cortical beta recordings as a reference signal. The two groups are composed of Parkinson’s patients undergoing deep brain stimulation surgery of either the STN or the GPi. Second, we investigate 150 Hz high-pass filtered LFP high-frequency activity (HFA) as a proxy of background multiunit activity (Meidahl et al., 2019; Moran and Bar-Gad, 2010), capturing fluctuations of the summed activity of a large number of cells and allowing us to define the co-modulation of our proxy for multiunit activity by cortical phase.

## Materials and methods

### Participants

We recorded 18 patients with idiopathic Parkinson’s disease (mean disease duration of 10 ± (STD) 5 years, mean age 62 ± 7 years, 16 males, also see **Table 1**) who underwent surgery for bilateral implantation of deep brain stimulation electrodes either to the STN (n = 11) or the GPi (n = 7). The STN and GPi cohort differed in age (GPi = 56 ± 4 years, STN = 65 ± 7 years, p = 0.006, t_16_ = −3.1) and symptom severity (but note the missing data points **in Table 1**, UPDRS ON: GPi = 29 ± 8, STN = 14 ± 10, Wilcoxon ranksum test: p = 0.017; UPDRS OFF: GPi = 60 ± 13, STN = 37 ± 14, p = 0.017, t_12_ = 2.8), but not in disease duration (p = 0.751) or daily levodopa equivalent dose (p = 0.816). Due to limited access to patients participating in postoperative recordings, we could not match the two groups.

**Table 1.** Patient details. Dom hand = dominant hand; Hand tested = hand used in the task, R = right; L = left; Med State = Order of the recorded medication state, UPDRS-III = Unified Parkinson’s disease rating scale part III ON/OFF levodopa (obtained not at the time of the recording but preoperatively), Disease dur = disease duration. L-Dopa dose (Levodopa equivalent dose) was calculated according to Tomlinson et al. (2010). Age and disease duration is provided in years.

| ID | Site | Dom. | Med state | UPDRS-III | Sex | Age | Disease |  | Electrode Model | L-Dopa dose |
| --- | --- | --- | --- | --- | --- | --- | --- | --- | --- | --- |
|  |  | Hand/Hand tested |  | ON/OFF |  |  | Dur |  |  |  |
| 1 | GPI | R/R | ON-OFF | 36/73 | m | 55 | 11 |  | Medtronic 3389™ | 940 mg |
| 2 | GPI | R/R | ON-OFF | 33/68 | m | 62 | 7 |  | Medtronic 3389™ | 2137.5 mg |
| 3 | GPI | R/R | ON | N/A | m | 50 | N/A |  | Medtronic 3389™ | N/A |
| 4 | GPI | R/R | OFF-ON | 17/49 | f | 52 | 11 |  | Medtronic 3389™ | 707 mg |
| 5 | GPI | R/R | OFF-ON | N/A | m | 59 | N/A |  | Medtronic 3389™ | N/A |
| 6 | GPI | R/R | OFF-ON | N/A | m | 58 | N/A |  | Medtronic 3389™ | N/A |
| 7 | GPI | R/R | ON-OFF | 28/48 | m | 58 | 8 |  | Boston Scientific DB-2202™ | 1513 mg |
| 8 | STN | L/L | OFF-ON | N/A | f | 72 | 24 |  | Boston Scientific DB-2201™ | 1900 mg |
| 9 | STN | R/L | ON | 38/63 | m | 62 | 11 |  | Medtronic 3389™ | 1014 mg |
| 10 | STN | R/L | ON | 26/53 | m | 69 | 9 |  | Medtronic 3389™ | 798 mg |
| 11 | STN | R/L | OFF-ON | 6/37 | m | 60 | 6 |  | Boston Scientific DB-2202™ | 2084 mg |
| 12 | STN | R/L | ON-OFF | 9/17 | m | 56 | 3 |  | Boston Scientific DB-2201™ | 328 mg |
| 13 | STN | R/R | ON-OFF | 10/31 | m | 75 | 11 |  | Medtronic 3389™ | 565 mg |
| 14 | STN | R/L | OFF-ON | 8/32 | m | 68 | 8 |  | Medtronic 3389™ | 720 mg |
| 15 | STN | R/R | ON | 9/49 | m | 54 | 14 |  | Medtronic 3389™ | 3063 mg |
| 16 | STN | R/L | ON | 10/25 | m | 64 | 13 |  | Medtronic 3389™ | Apomorph. pump |
| 17 | STN | R/R | ON | 13/31 | m | 67 | 3.5 |  | Medtronic 3389™ | 2173 mg |
| 18 | STN | R/R | On | 15/33 | m | 68 | 10 |  | Boston Scientific DB-2202™ | 1765 mg |

The surgery was performed 3-6 days prior to the recording. Recordings were enabled with DBS electrode extension cables, which were externalized through the scalp in the first operation and then connected to a subcutaneous DBS pacemaker in a second operation. Three different macroelectrode models were used: Model 3389™ from Medtronic Neurologic Division, Model DB-2201™ (cylindrical octopolar) or DB-2202™ (directional) from Boston Scientific. Surgeries and recordings were performed at one of the following three sites: King’s College Hospital, London, University College Hospital, London, or the John Radcliffe Hospital, Oxford, UK. The same data was published previously showing that task-related changes in the power of beta oscillations are broadly similar in the STN and the GPi (Fischer et al., 2019). The study was approved by the local ethics committees and patients provided written informant consent before taking part in this study.

### Experimental paradigm

#### Motor control task

The task was described previously (Fischer et al., 2019) and required patients to control the size of a box displayed on a computer screen by regulating the pressure of a pen on a graphic tablet (Wacom Intuos CTL-480, small). The task was visually guided by displaying a blue target box on a laptop screen, which randomly changed its size abruptly on average every 3.7s ± 2.2s. Patients were asked to manipulate the size of a black cursor box to match the blue target box by increasing or decreasing the force they applied with the pen on the tablet. If the blue box suddenly expanded, patients had to increase the force, and if it became smaller, they had to decrease it. The task was performed with the dominant hand, but if patients had severe tremor or rigidity in this hand, they used their non-dominant hand. All analyses were performed on the contralateral LFP and motor cortex activity as this is where the movement task should result in strongest engagement of neural activity. All patients were recorded ON medication, and a subset was also recorded OFF medication (5 STN, 6 GPi), which was analysed for a previous publication (Fischer et al., 2019). Because the sample size was strongly reduced for the STN cohort and thus a between-group comparison would be underpowered, we did not include an OFF medication comparison. Each continuous recording in the pen task lasted two minutes and patients performed four recordings with breaks in-between.

In half of all recordings, a second box coloured in red was displayed in addition to the blue target box, which randomly differed in size as distracting stimulus. Patients were instructed to ignore the red box and did so successfully. A demo of the task including the distractor box can be viewed on YouTube (https://www.youtube.com/watch?v=v7EbPiZB-dM [accessed April 2020]). As we were only interested in the task-related vs. resting dynamics in this study, we pooled the data across all pen task recordings irrespective of whether the red box was present.

#### Data recording

All LFP and EEG channels were recorded with a TMSi Porti amplifier (2048 Hz sampling frequency, with a 553 Hz anti-aliasing low-pass filter) with a common average reference. EEG electrodes were placed over (or close to if sutures had to be avoided) Fz, Cz, Pz, Oz, C3 and C4 according to the international 10–20 system. For one patient, the electrode over ipsilateral motor cortex was intermittently inactive and thus had to be excluded. The force exerted with the pen on the graphic tablet was recorded in a separate text-file but was not analysed in this study. Before patients performed the task, we also recorded two minutes of rest recordings.

### Analyses of EEG and LFP recordings

#### Data pre-processing

The data were down-sampled to 1000 Hz and re-referenced offline to make sure that the basal ganglia LFP and cortical EEG signals did not share any common reference signals (also described in Fischer et al., 2019). Briefly, the EEG electrode that was positioned over the region of the contralateral motor cortex (C3 or C4 depending on whether the task was performed with the right or left hand) was re-referenced to the average of all recorded EEG channels (Fz, Cz, Pz, Oz, C3 and C4). To obtain bipolar signals for the LFPs to exclude volume-conducted cortical activities (Marmor et al., 2017), the difference between the raw signal of two neighbouring DBS electrode contacts was computed. If single channels saturated or were inactive (6 of 36 electrodes), the remaining surrounding contacts were subtracted instead. The difference between the lowest and highest contacts was also included as additional bipolar configuration, as in some cases (8 of 36 electrodes), movement-related beta modulation across contiguous bipolar configurations was not clearly visible. Artefacts in the resulting re-referenced signals were excluded by visual inspection. Finally, for each patient, we pre-selected the bipolar configuration with the strongest movement-related beta modulation in the ON medication condition. To this end, we computed t-scores of 15-30 Hz beta power averaged across a −300:100ms window around the force adjustments, as reported previously (Fischer et al., 2019).

### Extracting the cortical beta phase and the high-frequency activity from the basal ganglia LFP

Phase and amplitude of EEG and LFP signals were obtained with the Hilbert transform. Before estimating the phase/amplitude, signals were pre-processed to isolate frequency bands of interest. In particular:

a. The phase of the EEG signal was obtained by band-pass filtering the data (Butterworth filter, filter order = 4, passed forwards and backwards, *fieldtrip*-functions *ft_preproc_lowpassfilter* and *ft_preproc_highpassfilter*) and extracting the phase of the analytic signal obtained with the Hilbert transform (using the *MATLAB* functions *angle(hilbert(filteredSignal))*). The EEG filter band was 15-30 Hz for all analyses apart from the plots in **Figure 5**, which show control analyses for 8-12 Hz, 30-40 Hz and two beta sub-bands (15-20 Hz, 20-30Hz). Our rationale for using the 15-30 Hz band as beta range of interest was that it showed clear movement-related modulation of motor cortical and subcortical activity in our previous publication and also significant cortico-subcortical phase coupling across all recordings (Fischer et al., 2019). The task-related modulation and significant coupling suggests a functional role of cortical 15-30 Hz oscillations, which we want to delineate further in this article.
b. The STN/GPi HFA was calculated by high-pass filtering the LFP recording with a cut-off frequency of 150 Hz (*fieldtrip* Butterworth filter, filter order = 4, passed forwards and backwards), taking the amplitude of the Hilbert-transformed signal, and subsequently smoothing the signal by low-pass filtering with a cut-off of 50 Hz to retain only the slower fluctuations in HFA amplitude. The cut-off of 150 Hz captures broadband activity between 150-550 Hz due to the 553 Hz anti-aliasing low-pass filter of the amplifier. Amplitude increases in this range correlate with firing activity (Moran and Bar-Gad, 2010; Winestone *et al*., 2012; Meidahl *et al*., 2019; although some controvery was reported previously: Yang *et al*., 2014; Storzer *et al*., 2015) and exclude potential oscillatory activity in the gamma range closer to 100 Hz, which shows distinct movement-related modulation (Litvak et al., 2012).

The co-modulation pattern of the HFA with the phase of cortical beta was computed to show if consistent phase-amplitude coupling existed between the cortical beta phase and STN/GPi HFA. It was plotted as HFA amplitude on the y-axis against the cortical beta phase on the x-axis (showing one full cycle from −*π* to + *π*, see **Figure 1B**). More specifically, it was obtained by subdividing the continuous beta phase signal into 63 bins (each tagging 0.1 rad wide periods) ranging from −*π* to + *π*. We only tagged periods where the EEG beta amplitude exceeded the 25^th^ percentile, to include only periods with a meaningful beta phase while still using most of the data to keep our signal-to-noise ratio high. In a next step, we computed the median of the HFA coinciding with each of the 63 different tagged time periods. The resulting curve for each patient was then smoothed (using the MATLAB function *smooth* over 20 phase bins, applying a moving average filter) to facilitate permutation-based statistical comparisons with cluster-based statistics that rely on detecting contiguous activity changes across subjects. Without smoothing, the modulation patterns across subjects would be more variable, making large clusters of significant points less likely. To normalize the HFA modulation pattern obtained for each patient, the relative HFA change was computed by subtracting the mean HFA (across all time points) from the original HFA pattern, dividing all values by the same mean and multiplying them by 100.

**Figure 1.**
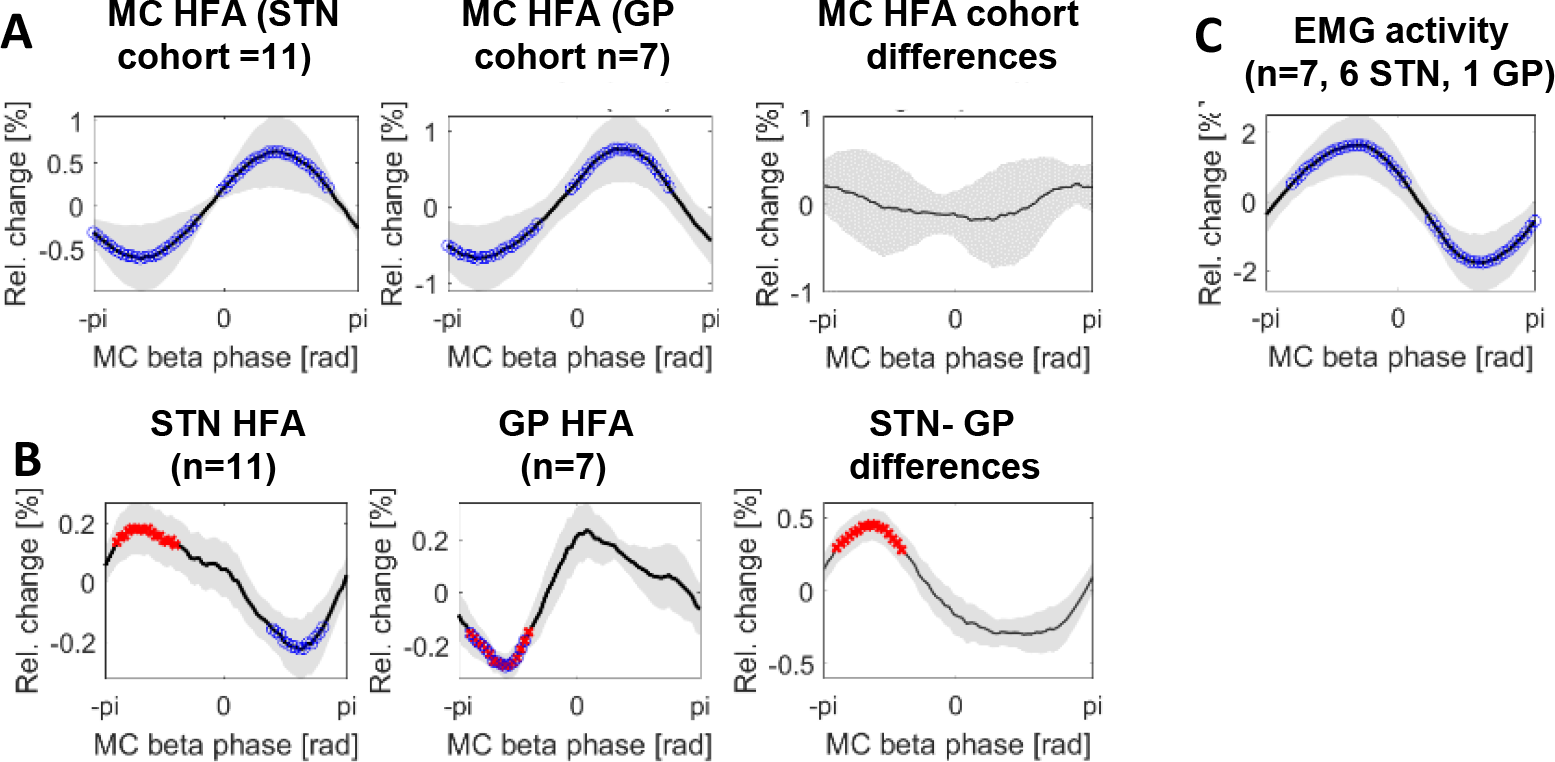
HFA modulation during the task. **A** The first two plots show the modulation of the motor cortical HFA relative to the motor cortical beta phase. Both signals were obtained from the same EEG electrode that was used to extract the phase of beta activity for all other analyses (plotted along the x-axis). They confirmed that local motor cortical HFA cofluctuated with the local beta phase similarly in both cohorts and show which beta phase coincided with reduced and increased cortical excitability as indexed by increases and decreases in HFA. Blue circles denote clusters that are significantly different from zero after cluster-based correction for multiple comparisons. Grey shaded areas show standard errors of the mean across patients. The third plot shows that the difference curve between the STN cohort (n=11 patients) and GPi cohort (n=7) was relatively flat, confirming that the HFA modulation was similar in the two groups. This was performed as control analysis to ensure that the polarity of cortical beta activity did not differ between the two cohorts. **B** STN HFA peaked close to −π while GPi activity peaked close to 0π. The points highlighted in red show where the STN and GPi modulation patterns were significantly different to each other. The 3^rd^ plot shows the difference between the two patterns. Note that both peaks and troughs for both STN and GPi modulation patterns would be significantly different from zero (and highlighted with blue circles) if no multiple comparison correction would be applied. **C** Electromyographic (EMG) activity of the forearm muscle also was co-modulated with the phase of cortical beta activity, showing the average across all patients, where EMG was recorded. If split up, the co-modulation pattern is highly similar in the STN (n=6) and the GPi (n=1) patient recordings.

Note that the high-pass filter used to extract the HFA successfully suppressed low-frequency fluctuations, as the HFA modulation pattern remained identical when the sign of the LFP bipolar signal was flipped before filtering (turning the peaks of beta oscillations into troughs).

### Analyses of motor cortical HFA and EMG

As a control analysis, we also investigated the pattern of local motor cortical HFA co-modulation with the beta phase recorded from the same electrode. This was to ensure that these patterns were similar in the STN and the GPi cohort (**Figure 1A**). As before, the HFA modulation pattern remained identical when the sign of the EEG signal was flipped before filtering. Note that it has been shown that high-frequency activity can indeed be captured with EEG (Freyer et al., 2009; Hashimoto, 2000), although the estimates are expected to be noisier than those extracted from LFPs because the signal is strongly attenuated when recorded from outside the skull (Buzsáki et al., 2012) and may be contaminated by scalp EMG. The procedure for computing this pattern was the same as described above for the LFP. In a subset of patients (n=7) we also recorded EMG activity of the extensor digitorum muscle (**Figure 1C**). A high-pass filter at 150 Hz would be too high for EMG as the frequency content of surface EMG discharges extends to lower frequencies. However, we still wanted to limit the effect of any movement artefact which might contaminate EMG and EEG. Accordingly, we filtered EMG recordings with a cut-off of 40 Hz, rectified them and low-pass filtered them with a cut-off of 50 Hz as above. The rest of the procedure again remained the same and the co-modulation pattern looked very similar with a cut-off of 150 Hz.

### Cross-correlation

We computed the STN and GPi HFA modulation patterns not only on the pen task data but also on a separate rest recording. To evaluate their similarity and examine if any systematic temporal shifts existed between the two modulation patterns, we computed cross-correlations (using the MATLAB function *xcorr*) between the two HFA modulation patterns obtained from the task and rest recording for each subject and then plotted the average cross-correlations across subjects.

### Power spectra

The power spectral density between 5-40 Hz and 150-500 Hz was estimated using Welch’s method (MATLAB function pwelch, 4s hamming window, 50% overlap). The method attenuates noise by computing the discrete Fourier transform in overlapping windows and averaging across the resulting spectra. Before plotting the average across patients, all spectra were normalized by dividing them by the mean of 5-500 Hz power averaged across rest and task recordings (excluding line noise peaks at 50 ±5 Hz and multiples of 50 Hz).

### Statistical analyses

Cluster-based permutation statistics were computed for all significance tests that were performed across multiple frequency or time bins. Hence, to test whether EEG-to-LFP HFA coherence or the HFA modulation patterns were significant, we created 500 surrogate data sets making up our null-hypothesis distribution. Each surrogate data set was created by cutting the vector containing the LFP HFA of each recording at a random point (drawn from a uniform distribution ranging from 0s to the end of the recording) and swapping the second part with the first part, thus destroying the original relationship between the LFP HFA vector and the EEG beta fluctuations while retaining the rhythmic structure within the HFA and EEG recordings. The scalar coherence values (for the frequencies ranging from 5-40 Hz) or amplitude values (for all 63 phase bins) for each surrogate data set were then computed exactly as described above for the original data. P-values for each frequency or phase bin were then computed by counting how many of the absolute surrogate coherence/amplitude patient-averaged values (*V_p_*) were larger than or equal to the absolute original patient-averaged values (*V_orig_*) and dividing the outcome by the number of permutations (*N_p_* = 500).

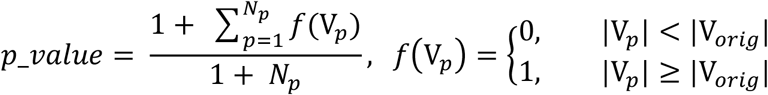

The logic here is that the shuffled surrogate data sets should result in lower coherence values (or in smaller amplitude peaks and troughs for the modulation pattern) than the original data, if the original effects are significant. The number 1 was added to both the nominator and the denominator to avoid p-values of 0 and to factor in that the exact p-value should be at least 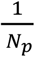 (see section 4.2 from Ernst, 2004). The same procedure is done also for each of the surrogate data sets (using the respective *V_p_* instead of *V_orig_*) resulting in p-values for each surrogate data set. For example, when testing if the HFA amplitude modulation was significant, 63 p-values (one for each phase bin) were computed. If any phase bins had a p-value below 0.05, the sum of all amplitude values within a contiguous cluster of significant phase bins (or frequency bins in the case of testing for significant coherence) was stored. If several significant clusters existed in one surrogate data set, then only the largest sum was stored to serve as a comparison value for all clusters found in the original data, which is key for keeping the false positive rate at 0.05 (Maris and Oostenveld, 2007). The final step to compute the p-values for each original cluster of significant phase/frequency bins was to test how many of the surrogate cluster sums were larger than or equal to the original sum and again divide by the number of permutations.

To evaluate differences in the power spectra between the task and rest recordings, a similar principle was used: Instead of cutting and swapping the temporal sequence, which would have not made a difference for the power estimate, we shuffled the task labels randomly for each patient 500 times to compute 500 surrogate differences and calculate cluster-based corrected p-values as described above.

## Results

### Co-modulation of STN and GPi high-frequency activity with motor cortical beta during task

As control analysis, we tested first if motor cortical HFA co-fluctuated with the beta phase measured from the same EEG electrode and if the modulation pattern was the same in both groups. Both groups showed a highly similar pattern of motor cortical HFA modulation relative to the local beta phase (**Figure 1A**), confirming that the EEG beta phase provided a consistent measure of motor cortical beta activity that did not differ between the two groups. In contrast to the similarity of motor cortical HFA, the HFA modulation patterns obtained from the STN and the GPi were markedly different (**Figure 1B**): group average STN activity peaked close to −*π* while GPi activity peaked at around 0*π*. The peak in STN activity coincided with the trough of GPi activity, showing opposite patterns, resulting in significantly different HFA amplitudes between the two groups (highlighted in red).

Finally, we also investigated the modulation of EMG activity (**Figure 1C**, n = 6 STN, n = 1 GPi). The group average peak was shifted by about half a cycle relative to the peak of motor cortical HFA.

### Is the HFA modulation task-specific?

All patients were also recorded at rest for two minutes. GPi HFA during rest evolved similarly as during the task (**Figure 2A**, GPi ON: peak p = 0.016, trough p = 0.032). To test more formally, how similar the modulation was between the rest and task condition, we computed cross-correlations (**Figure 2B**). If the cross-correlation has its positive peak at 0 and its negative maximum at −pi or pi, then the rest and task modulation patterns were similar without any temporal shift between each other. This was roughly the case for GPi HFA, the peak was shifted only slightly to the right, indicating a slight shift of only a few milliseconds between the average modulation patterns recorded during rest and task.

**Figure 2.**
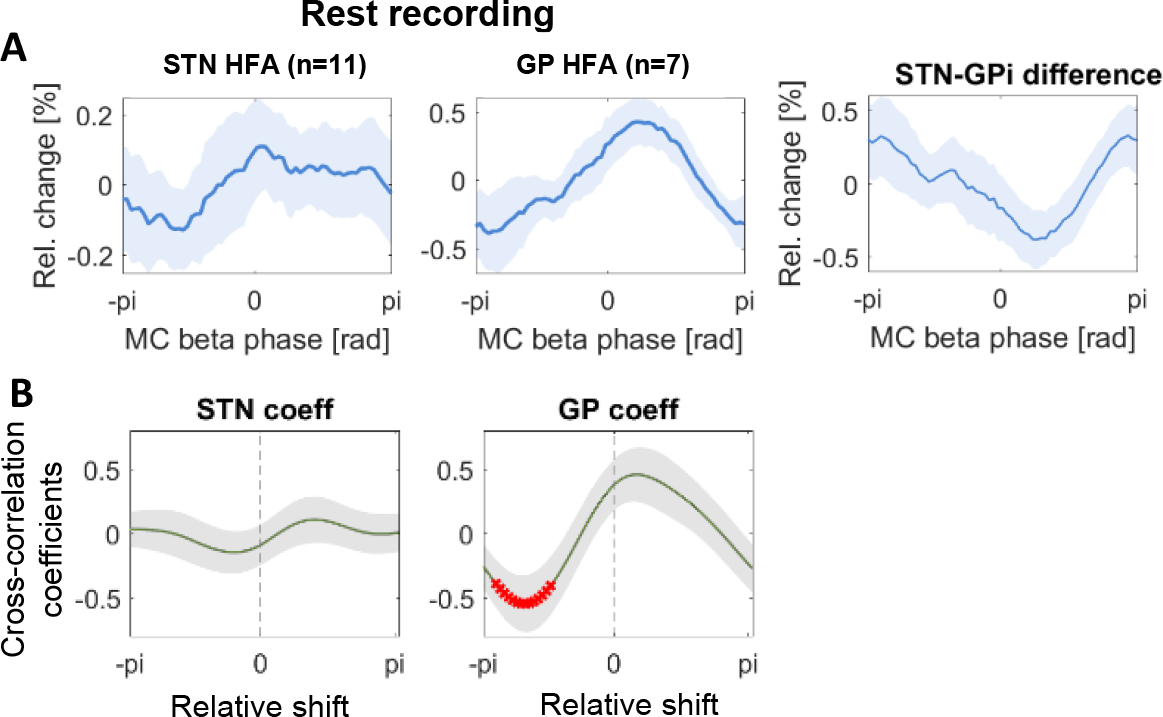
HFA modulation when patients were at rest. **A** Modulation patterns were computed as in Figure 2B, but on the rest recordings instead of the task recordings (with the data obtained form the same contact pairs). **B** Average of correlation coefficients across patients, which were obtained by crosscorrelating the two average modulation patterns obtained during task and rest for each patient. The correlation coefficient at 0 shows the correlation between the two patterns with a temporal shift of 0. HFA patterns in the STN were not consistently similar (low correlation at 0 pi) or temporally shifted relative to each other, as the cross-correlation pattern showed no distinct peak or trough. Conversely, GPi modulation patterns were similar during task and rest, which is shown by a positive correlation at around 0 rad and a negative correlation at around −pi rad (compare also the GPi pattern of Figure 3A with Figure 2B). Red points show where a cluster of correlation coefficients was significantly different from zero.

In contrast, the STN HFA modulation pattern was more variable during rest compared to the task-related modulation as shown by the increased standard error across the eleven patients in **Figure 2A** and the lack of significant modulation (peak p= 0.449, trough p = 0.333). The cross-correlation based on the STN recordings showed no significant peaks (**Figure 2B**).

To test if the increased variability may be due to the reduced recording length of the resting data, we computed the task-related HFA on shorter segments taken from the start (**Figure 3A**) and the end (**Figure 3B**) of the motor control task. This showed that data length alone was unlikely to account for the difference in STN HFA modulation from that recorded during task as the control analysis was able to capture consistent modulation patterns in the shorter task recordings for both cohorts. Significant clusters in the modulation patterns were only present in the segment taken from the end and resemble the clusters in the full-length recording (with both trough and peak now being significantly different). Consequently, we tested if the absence and presence of significant clusters in the two segments was related to a difference in motor cortical or subcortical beta power between the two segments. This was tested for all recordings and also separately for the STN and GPi recordings, but no significant differences were found. Additionally, we compared the motor cortical EEG and basal ganglia power spectra between the rest and the task condition to investigate any changes in power, which could have been linked to changes in HFA modulation (**Figure 4**). We found no significant differences in the beta range or of power between 150 and 500 Hz. The only significant difference was increased 8-12 Hz motor cortical EEG activity in the STN cohort when being at rest (**Figure 4A**, red shaded area).

**Figure 3.**
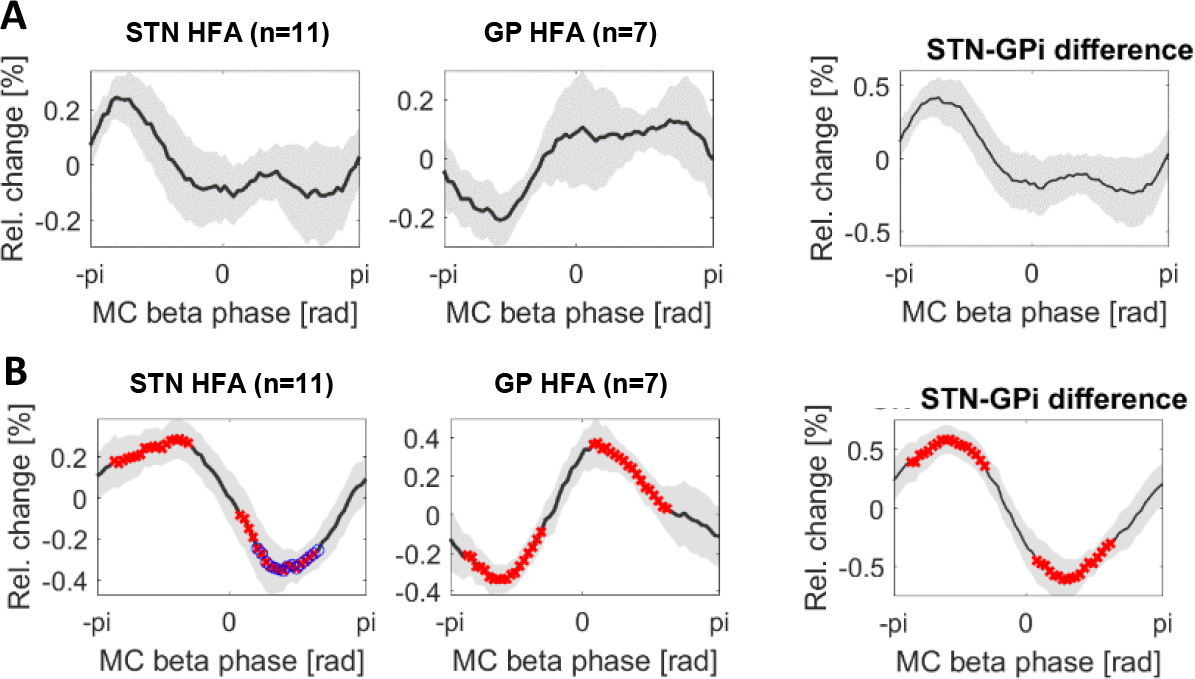
HFA modulation during the task: Control analyses on shorter segments. A+B show the same analysis as Figure 2B but on shorter segments of the task recording matched in length to the rest recording. The segments for (A) were taken from the beginning of the task recording, the segments for (B) were taken from the end. Note that both patterns are very similar as in Figure 2B, but that the differences were only significant in B, suggesting that although the patterns likely are meaningful, the lack of significance in Figure 3 may be due to the shorter recording segments, increasing the variability of the average pattern across patients.

**Figure 4.**
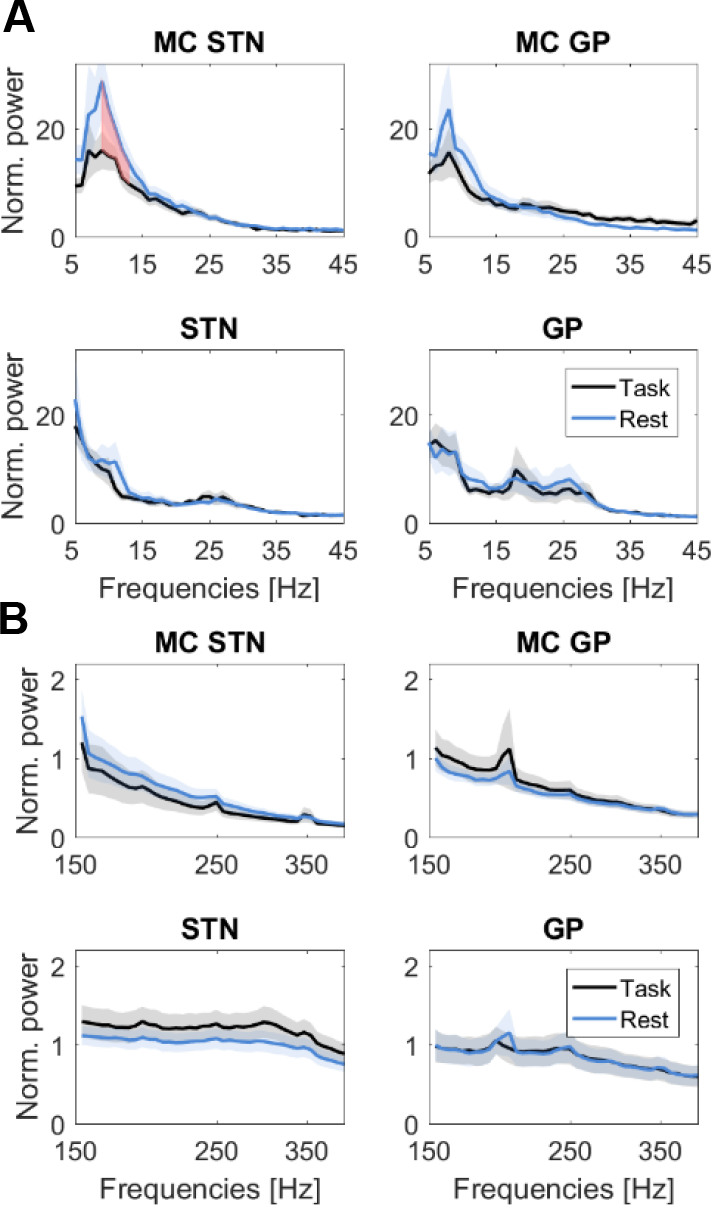
Power spectra. **A** Power spectra between 5-45 Hz of the motor cortical (MC) EEG signals (1^st^ row) for the STN (n = 11) and GPi cohort (n = 7) and for the STN and GPi LFP signals (2^nd^ row) during task (black line) and rest (blue line). The red shaded area shows that 8-12 Hz motor cortical alpha power was significantly higher during rest in the STN cohort, but no other differences in beta power were significant. **B** Power between 150 and 500 Hz was not significantly different between task and rest. Peaks at 200, 250 and 300 Hz are due to 50 Hz line noise. Because line noise is stable, its constant amplitude does not affect the HFA modulation patterns.

### Is the HFA modulation frequency-specific?

To test if the task-related modulation of HFA is frequency-specific, we computed the modulation pattern for four different frequency bands: 8-12 Hz, 15-20 Hz, 20-30 Hz and 30-40 Hz. We saw no significant HFA modulation or difference in HFA modulation when calculating it based on the phase of cortical alpha activity (**Figure 5**; 8-12 Hz: STN peak p = 0.080, trough p = 0.100; GPi peak p = 0.503, trough p = 0.535) or for cortical 30-40 Hz activity (STN peak p = 0.425, trough p = 0.501, GPi peak p = 0.259, trough p = 0.281). We also subdivided the 15-30 Hz band into low (15-20 Hz) and high beta (20-30 Hz). The modulation patterns were highly similar and the difference between the STN and GPi patterns were the same. The blue clusters highlighting significant HFA amplitude deviations from zero, were present only in the 15-20 Hz band for the STN and in the 20-30 Hz band for the GPi, indicating stronger across-subjects consistency for the respective bands. In addition, the p-values for the peaks and troughs showed that HFA amplitude was significantly modulated in both nuclei (15-20 Hz: STN peak p = 0.018, trough p = 0.002, GPi peak p = 0.012, trough p = 0.018; 20-30 Hz: STN peak p = 0.090, trough p = 0.026, GPi peak p = 0.006, trough p = 0.004), highlighting again the similarity.

**Figure 5.**
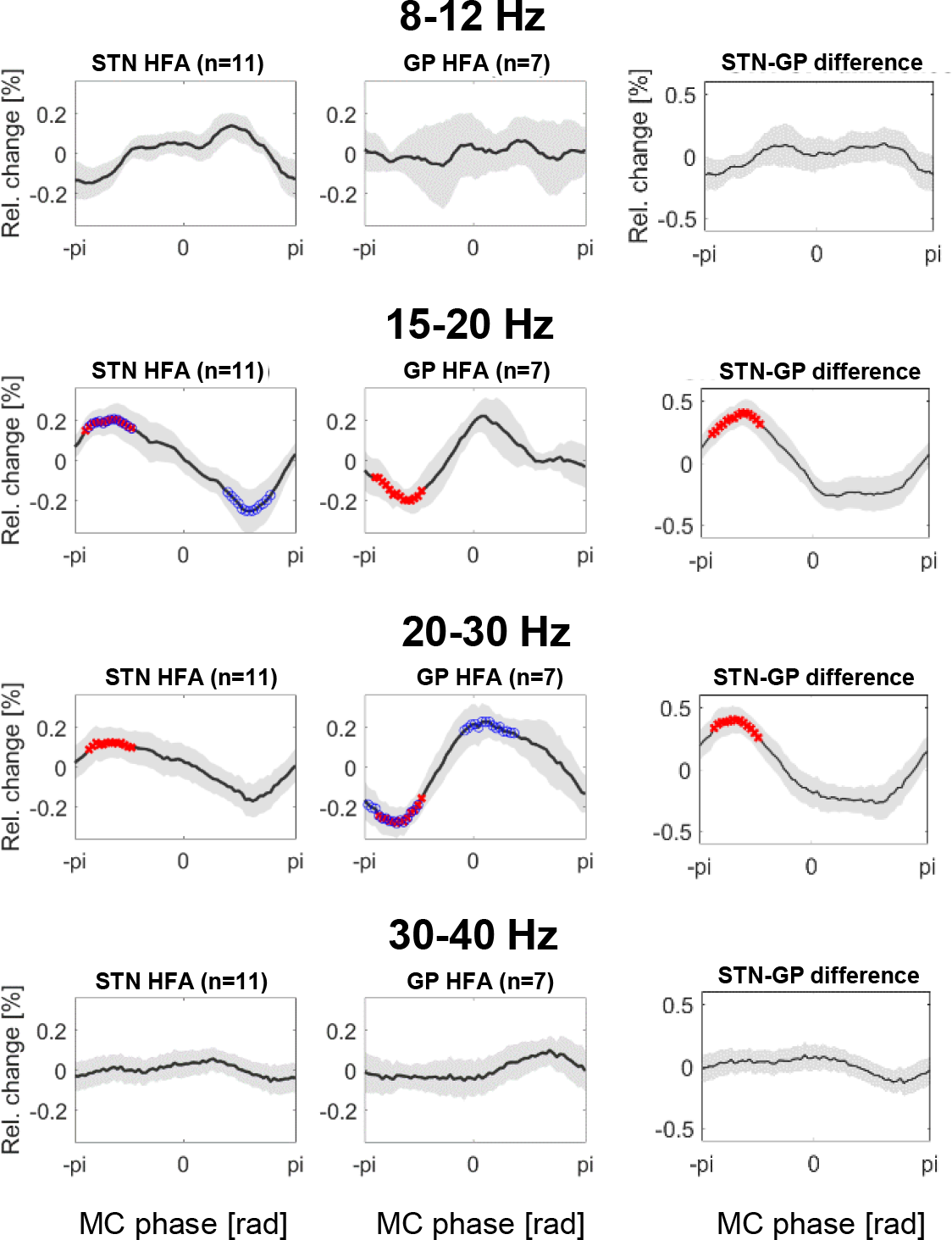
Frequency-specificity of HFA modulation. The individual rows show the different HFA modulation patterns originating from filtering the motor cortical EEG signal in four different frequency bands before extracting the phase and computing how the basal ganglia HFA was co-modulated relative to it. Significant modulation was only present in the two beta sub-bands (15-20 Hz and 20-30 Hz).

## Discussion

We found that during performance of a low-force motor control task, HFA recorded from the STN and GPi was significantly co-modulated with the phase of motor cortical beta activity and that their modulation patterns were inverted relative to each other. The relatively large temporal shift of half a beta cycle implies that rhythmic cortical firing during task-related beta oscillations (Baker et al., 1999; Canolty et al., 2012; Denker et al., 2011; Donoghue et al., 1998; Murthy and Fetz, 1996) is not simply transferred to the GPi via monosynaptic STN connections, because the difference of about half a cycle translates to a delay of about 25ms (for a beta frequency of 20 Hz). This difference dissipated when recordings were made at rest, which was due to a state-dependent diminution of STN but not GPi HFA modulation. Whatever causes GPi activity to co-fluctuate with cortical beta activity thus likely involves additional parts of the network, such as GPe or striatal activity (Shink and Smith, 1995; Singh and Papa, 2019; Smith et al., 1994).

### Functional relevance of cortico-subcortical coupling

The comparison between task-related and resting activity showed that GPi activity was consistently co-modulated with motor cortical beta activity, irrespective of whether patients performed the task. STN activity instead was not consistently entrained to the phase of cortical beta when patients were at rest – hence, only task-engagement appeared to result in significant entrainment of STN activity. This resembles previous reports based on rest recordings, where no such STN-MC coupling was found (van Wijk et al., 2016). It is also in line with the observation that cortico-STN beta coherence is increased during sustained contractions (Hirschmann et al., 2013; Marsden et al., 2001), while it is decreased during phasic movements (Lalo, 2008).

We would like to acknowledge that the rest recordings were shorter than the task recordings (2 min vs. 8 min). Thus, the lack of consistent coupling in the STN may be due to the reduced data length. However, reducing the length of the task data, such that it was matched to the rest recording, still showed similar task-related HFA patterns as when calculating it based on the full-length data. Note that the shape of the STN HFA resting pattern was very different to the task-related pattern, highlighting again that STN HFA was not engaged in the same way as when patients performed the motor control task.

Recent studies have shown that cortical beta oscillations can appear and fade away independent of changes in firing rates (Confais et al., 2019; Rule et al., 2017) and thus could be a flexible and energy-efficient way to regulate the patterning of downstream activity if the recipient sites are not hijacked by pathological activity. Increased beta-HFA phase-amplitude coupling (PAC) in Parkinson’s disease due to dopamine depletion has been found locally in motor cortex, the STN and the GPi (de Hemptinne et al., 2013; Lopez-Azcarate et al., 2010; Swann et al., 2015; Tsiokos et al., 2017; van Wijk et al., 2016). The strength of such local coupling correlates with symptom severity (Lopez-Azcarate et al., 2010; Özkurt et al., 2011; Rajagopalan et al., 2019) and is stronger contralateral to the more severely affected side (Shreve et al., 2017), which may reflect compromised flexibility of network dynamics. However, intake of dopaminergic medication also resulted in increased coupling between the phase of STN high-beta activity and local HFA in a 300-400 Hz band and was associated with greater motor improvement (Ozturk et al., 2019), suggesting that some forms of beta-HFA PAC can even be beneficial for movement control.

Our study showed that STN/GPi HFA was coupled to the motor cortical beta phase while patients engaged in our motor control task. The task involved sustained isometric contractions, maintaining a visually-cued force level. In previously published analyses of the same data, we have shown that force adjustments were finished more accurately when motor cortical and basal ganglia beta oscillations were relatively high (Fischer et al., 2019). Hence, increased mortor cortex beta-basal ganglia HFA PAC may even benefit the process of controlling sustained contractions. Our approach will also be suitable to investigate how HFA modulation patterns change during dopamine withdrawal, which may play a role in the emergence of motor impairments, perhaps even before the appearance of excessive local synchrony (Degos et al., 2009; Devergnas et al., 2019; Leblois et al., 2007).

### Future implications

One key novel aspect compared to previous PAC studies, which tend to focus on summary statistics of coupling strength but not modulation patterns, is that we showed consistent modulation patterns across the STN and GPi recordings.

Studies that aim to understand how firing activity (or a proxy-measure as used here) is coupled to oscillations in distant but connected sites will be critical for understanding how shifts in the balance between functional network connectivities may change to fulfill different motor control demands. Ballistic movements, for example, seem to engage different beta dynamics compared to tonic contractions (Hirschmann et al., 2013; Marsden et al., 2001). The post-movement beta rebound is another example: It has been linked to feedback processing (Cao and Hu, 2016; Tan et al., 2016; Torrecillos et al., 2015), and recently we found that STN cells that are coupled to the post-movement rebound can also be coupled to motor cortical gamma oscillations at movement onset (Fischer et al., 2020), suggesting that distinct states of cortico-STN coupling are linked to distinct behavioural states. As our task involved continuous force regulation, our recordings did not allow us to assess HFA modulation relative to post-movement beta rebound activity, however, it will be interesting to see in future studies if the modulation patterns are the same as during isometric contractions.

### Choice of frequency bands

We chose to focus on 15-30 Hz as beta band of interest, because it showed task-related modulation in our previous publication (Fischer et al., 2019) despite the absence of a clear beta peak in the motor cortical power spectra. Not seeing a clear peak in a power spectrum does not necessarily mean that phase coupling cannot be present (Brunet et al., 2014), thus choosing a frequency band based on movement-related modulation seems to be better suited to defining a functionally relevant band. We have also shown that the modulation patterns are specific to the beta band as no significant modulation was present in the bands above or below (8-12 and 30-40 Hz). In addition, our analyses of the low- and high-beta sub-bands (15-20 and 20-30 Hz) produced similar modulation patterns, hence we could not gain any novel insights into the different nature of these bands, although differences in HFA coupling with these bands have been reported previously (Ozturk et al., 2019).

The frequency for HFA bands (in some publications also called “HFO” for high-frequency oscillations) depends on where PAC is computed: For motor cortical PAC, a range between 50-150 or 200 Hz seems appropriate (de Hemptinne et al., 2013; Swann et al., 2015), whereas subcortical HFA usually starts at 150 Hz ranging up to 400 Hz (de Hemptinne et al., 2013; Lopez-Azcarate et al., 2010; Meidahl et al., 2019; Tsiokos et al., 2017; van Wijk et al., 2016). PAC analyses within the STN showed significant coupling only for HFA above 150 Hz but not below (de Hemptinne et al., 2013; Yang et al., 2014). We used 150 Hz as cut-off for our analyses aiming to capture activity that co-fluctuates with the local beta phase, which in turn is coupled to motor cortical beta (Fischer et al., 2019). The broad band high frequency activities between 150 and 500 Hz (which is the cut-off frequency of the anti-aliasing filter of the amplifier) were quantified here as HFA in this study as no distinct peaks were present in the high frequency power spectra, and evidence suggests that multiunit activity may be represented in the background activity of DBS electrodes up to several 1000Hz (Winestone et al., 2012).

Finally, we did not compute co-modulation plots of subcortical HFA relative to the phase of local beta, because we were primarily interested in the consistency of modulation patterns across recordings, and the polarity of local beta derived from DBS electrodes would need to be standardized before computing a group average. This is because the polarity depends on the order of subtraction of the two signals used to calculate the bipolar recording. Flipping the order depending on the HFA modulation pattern could standardize the polarity of bipolar signals (Fischer et al., 2020), but would automatically result in a consistent alignment of the peaks of the HFA modulation patterns across recordings, resulting necessarily in a significant modulation pattern due to the circular procedure.

### Limitations

One caveat we would like to acknowledge is that we may have also captured GPe activity with the GPi recordings considering that the GPi and GPe are located directly next to each other (Parent and Hazrati, 1995). Clinical measures confirmed the intended targeting in the form of responsiveness to stimulation and post-operative imaging, however, the diverse imaging platforms and procedures used across the three surgical centres precluded precise localization at the group level. This presents a major limitation, and the exact patterns and directionality of how activity in defined basal ganglia nuclei co-fluctuates relative to cortical beta should be followed-up in non-human primate recordings, or in patient cohorts in whom standardised, high resolution imaging is available.

Another caveat of our between-group comparison is that the target area of the DBS surgery generally is chosen according to the patient’s disease profile – our GPi cohort tended to be more affected by dyskinesia prior to the surgery, in keeping with the fact that prominent dyskinesias are often cited as an indication for GPi target selection (Liu et al., 2019; Ramirez-Zamora and Ostrem, 2018). This raises the possibility that pathological network activity may have differed to some extent between our GPi and STN patient cohorts. Moreover, the GPi cohort was significantly younger and had more pronounced motor symptoms. Two STN patients had some tremor during the recordings but performed the task with the non-affected hand, and two GPi patients had mild dyskinesia during the task.

Finally, we would like to acknowledge that our study is limited in that recordings took place only a few days after the DBS surgery, when micro-lesions caused by the electrode insertion procedure may alter activity.

### Conclusions

In summary, we have demonstrated that high-frequency (>150 Hz) activity recorded from DBS electrodes and its coupling to cortical oscillations recorded with EEG can serve as highly informative signal about how basal ganglia activity can be differentially modulated in different sites. Activity in the STN and the GPi was entrained to the phase of motor cortical beta activity while patients performed an isometric force control task. Their modulation patterns were inverted, suggesting that GPi activity fluctuations are not simply driven by glutamatergic STN afferents. Instead, other inputs, such as GABAergic inhibition from the striatum or the GPe, must play a significant role in crafting the GPi activity fluctuations, at least in the ON medication state. In contrast to GPi modulation, STN modulation was less pronounced during rest, suggesting disengagement of STN HFA from cortical beta oscillations in this state. Our findings should be considered preliminary, as our GPi cohort was small, the clinical characteristics of the STN and GPi cohorts were not matched and detailed imaging is unavailable. Nevertheless, visualizations of HFA co-modulation patterns as shown in this study may help define the impact of beta synchronization on activity at different sites of the cortico-basal ganglia-thalamo-cortical network and may help shed light on state-dependent changes in network dynamics.

## Acknowledgements

This work was supported by the Medical Research Council [MC_UU_12024/1], Rosetrees Trust, EU [FP7-ICT 610391] and the National Institute of Health Research (NIHR) Oxford Biomedical Research Centre (BRC). HT was additionally funded by the Medical Research Council [MR/P012272/1]. The Unit of Functional Neurosurgery was supported by the Parkinson Appeal UK, and the Monument Trust. LZ is supported by the National Institute for Health Research University College London Hospitals Biomedical Research Centre.

